# Robustness and the evolution of length control strategies in the type III secretion system and flagellar hook

**DOI:** 10.1101/766733

**Authors:** Maulik K. Nariya, Abhishek Mallela, Jack J. Shi, Eric J. Deeds

**Author notes:** Laboratory of Systems Pharmacology, Harvard Medical School, Boston, MA USA. Department of Mathematics, University of California Davis, Davis, CA, USA.

## Abstract

Bacteria construct many structures, like the flagellar hook and the type III secretion system, that aid in crucial processes such as locomotion and pathogenesis. Experimental work has suggested two competing mechanisms bacteria could use to regulate length in these structures: the “ruler” mechanism and the “substrate switching” mechanism. In this work, we constructed a mathematical model of length control based on the ruler mechanism, and found that the predictions of this model are consistent with experimental data not just for the scaling of the average length with the ruler protein length, but also the variance. Interestingly, we found that the ruler mechanism allows for the evolution of needles with large average lengths without the concomitant large increase in variance that occurs in the substrate switching mechanism. These findings shed new light on the trade-offs that may have lead to the evolution of different length control mechanisms in different bacterial species.

## Length regulation in bacteria

Bacterial cells build a variety of structures on their exterior, *e.g*. flagella, pili, the Type III Secretion System (T3SS), etc., which help to carry out important functions such as locomotion, DNA transfer and pathogenesis [1]. In order to ensure effective function and to optimize the efficiency of transport, bacteria need to control the length of these structures with high precision. This poses a natural question about the regulation of the assembly process: how does the bacterial cell “know” when to stop growing these structures, especially since the structures themselves are outside the cell? It is as if one was trying to construct a building that is precisely thirty storeys high, from underground, with no way of directly looking at the structure as it is being built.

The solution to this fundamental problem has been perhaps best studied experimentally in two homologous model systems in gram-negative bacteria: the T3SS injectisome and the flagellar hook [1, 2]. The flagellum is an organelle that consists of a curved tubular structure known as the hook with one of its ends attached to a motor complex that terminates in the outer membrane of the cell and the other to a long flexible filament that “whips” as the motor turns the hook in order to provide locomotion. Even though it is the filament that facilitates the actual motion, the formation of the hook is a crucial step in the assembly process [2]. The T3SS injectisome needle is homologous to the flagellar hook; it consists of a narrow needle that grows in the extracellular region and helps transport effector proteins into the host cells during pathogenesis [3–6]. Morphologically, both T3SS injectisome and flagellum consist of a basal body that spans the inner and the outer membranes of the bacterial cell and has a narrow passage for the secretion of proteins. This basal body, which is often refered to as the base, along with a number of regulatory genes, controls the assembly of these structures [1]. Several mechanisms have been proposed to explain length control for these systems, and among these the two most popular ones are the substrate switching mechanism and the ruler protein mechanism.

In the substrate switching mechanism, there is an inner-rod that spans the inner and outer membranes of the cell and is located inside the base. Initially the inner-rod and the needle are assembled at the same time. Once the inner-rod is completed, the base stops growing the needle and switches substrates, secreting tip proteins to form a mature injectosome. In *Salmonella*, PrgJ has been identified to be the protein that forms the inner-rod, and Marlovits *et. al.* obsereved that overexpressing PrgJ resulted in shorter needles as compared to ones in wild type, while deleting PrgJ resulted in extremely long needles [7–9]. According to the ruler mechanism, there is a dedicated protein that acts a “molecular ruler.” During the assembly of the needle, the base periodically secretes a ruler protein and once the needle reaches an optimum length, the C-terminus of the ruler protein interacts with the base, signaling the base to stop growing the needle. Such ruler proteins have been identified not only in the T3SS (*e.g*. YscP in *Yersinia*) but also in the flagellar hook (*e.g*. FliK in *Salmonella*) [10, 11]. Furthermore, there is recent experimental evidence for the length control via the ruler mechanism in *Salmonella* injectisome, leading to an ongoing controversy about the functions of some of the proteins involved in assembly [12]. Experimental evidence for the ruler mechanism has primarily been provided by demonstrating that the average length of the needles/hooks increases linearly with increases in the length of the ruler protein itself. While there is thus qualitative agreement between available experimental data and these two alternative mechanisms, until recently very little had been done to quantitatively model these length control processes. This represents a fundamental problem for the field, because it is difficult to make rigorous, experimentally-testable predictions in the absence of a quantitative modeling framework.

In previous work, we constructed a mathematical model for the substrate switching mechanism. One of the key predictions of our model was that variance in the needle lengths depends quadratically on the average length [13]. We found this relationship to be independent of all but one underlying parameter of the model, which is the number of inner proteins required to complete the inner-rod. By comparing the our results with the available experimental data, we predicted the number of inner proteins subunits for completing the inner-rod to be around six, which is consistent with recent experimental work on determination of stoichiometry of PrgJ proteins in *Salmonella* [14]. The quadratic relation between the variance and average length implies large variability in lengths, especially for longer needles. This makes substrate switching a potentially inefficient mechinism for growing longer structures. Furthermore, it is possible that different bacterial species have evolved distinct mechanisms for length regulation. The only way to test such hypotheses is by understanding the ruler mechanism in a similar quantitative framework, and to compare and constrast the implications of the mathematical model with experimental data. While there is a rough agreement between needle length distributions and a simplified mathematical model of the ruler mechanism [15], to date there has been no quantitative exploration of how crucial parameters such as ruler length and the copy number of ruler proteins affect the average and variance in lengths.

In this work we developed a quantitative model for the ruler mechanism. Our initial model, in which the measurement of these structures takes place deterministically with the help of an exact ruler, predicts that variance should be *independent* of needle length and depend only on number of ruler and needle proteins in the bacterial cell. Our analysis of experimental studies in which the authors obtained length distributions of these structures for a wide range of ruler lengths revealed a linear relatiosnhip between the variance and the average length [10–12]. A more realistic approximation of our model, where we introduce uncertainty in the measurement process by the ruler protein, results in a linear relationship between the variance in needles and the average needle lengths if the average length is increased by simply inserting additional amino acid residues to the ruler protein, which is what is done experimentally. Interestingly, if we allow the needle protein length to co-vary along with other parameters (such as the ruler protein concentration), it is possible to increase the needle length *without* increasing the variance, which is likely impossible in the substrate switching mechanism. Taken together, our findings suggest an interesting set of evolutionary trade-offs between these two mechanisms: the substrate switching mechanism allows for energy-efficient control of the length of smaller structures, while the ruler protein mechanism allows for robust control of the lengths of longer structures at a higher energetic cost. Our results also suggest a very straightforward experimental program for further investigating the length control mechanism in a given bacterial species.

## Results

### Quantitative model for ruler mechanism

The assembly of macromolecular structures such as the flagellum and T3SS injectisome is a highly regulated process and involves interaction between a number of proteins [1, 2, 16]. Evidence suggests that there are dedicated ruler proteins, such as YscP in *Yersinia* and its homologue FliK in *Salmonella*, that are secreted during the assembly process. When the injectisome or the flagellar hook achieves an appropriate length, the C-terminal domain of the ruler protein interacts with the gate protein (FlhB in *Salmonella* and YscU in *Yersinia*) [10, 17, 18]. It is believed that this interaction causes the substrate specificity to switch, thus preventing the needle from growing any further [11]. To obtain a quantitative description for the ruler model we first constructed a mathematical model that considers the dynamical interplay between the base complex, the needle (or the hook) subunits and a molecular ruler of fixed length.

Figure 1 depicts the mechanism of the needle growth in T3SS injectisome using ruler proteins as per our intial mathematical model. Note that the figure only refers to the T3SS needle, but all the results described below are valid for the flagellar hook as well. Imagine a base being produced at time *t* = 0. During the assembly process, the ruler proteins and the needle proteins are secreted one at a time. The needle proteins polymerize in the extracellular region of the cell, resulting in growth of the needle itself. The needle length is represented by *L*, which is the distance between the surface of the outer membrane and the tip of the needle. At some time *τ*, after a sufficient number of needle protein bindings, the needle achieves a length that is suitable for the switch in substrate specificity. In our model, we denote the effective length of the ruler protein by *l*_*R*_, and when the needle reaches an appropriate length this facilitates the interaction between the ruler protein and the gate protein, switching the substrate specificity. Since the needle length has to be greater then or equal to the ruler length for maturation, all the ruler protein binding events that occur prior to the time *τ* are simply generating the necessary condition for maturation. As shown in the figure, the next ruler protein binding event occurs at some (random) later time *t*, and it is at this time that the needle stops growing and the base becomes “mature.” Since the maturation event occurs only when *L* ≥ *l*_*R*_, followed by a ruler protein checking event, for simplicity we ignored any uncertainties involved in the measurement process in our initial model, and as such we call it the *exact* ruler model.

**Figure 1:**
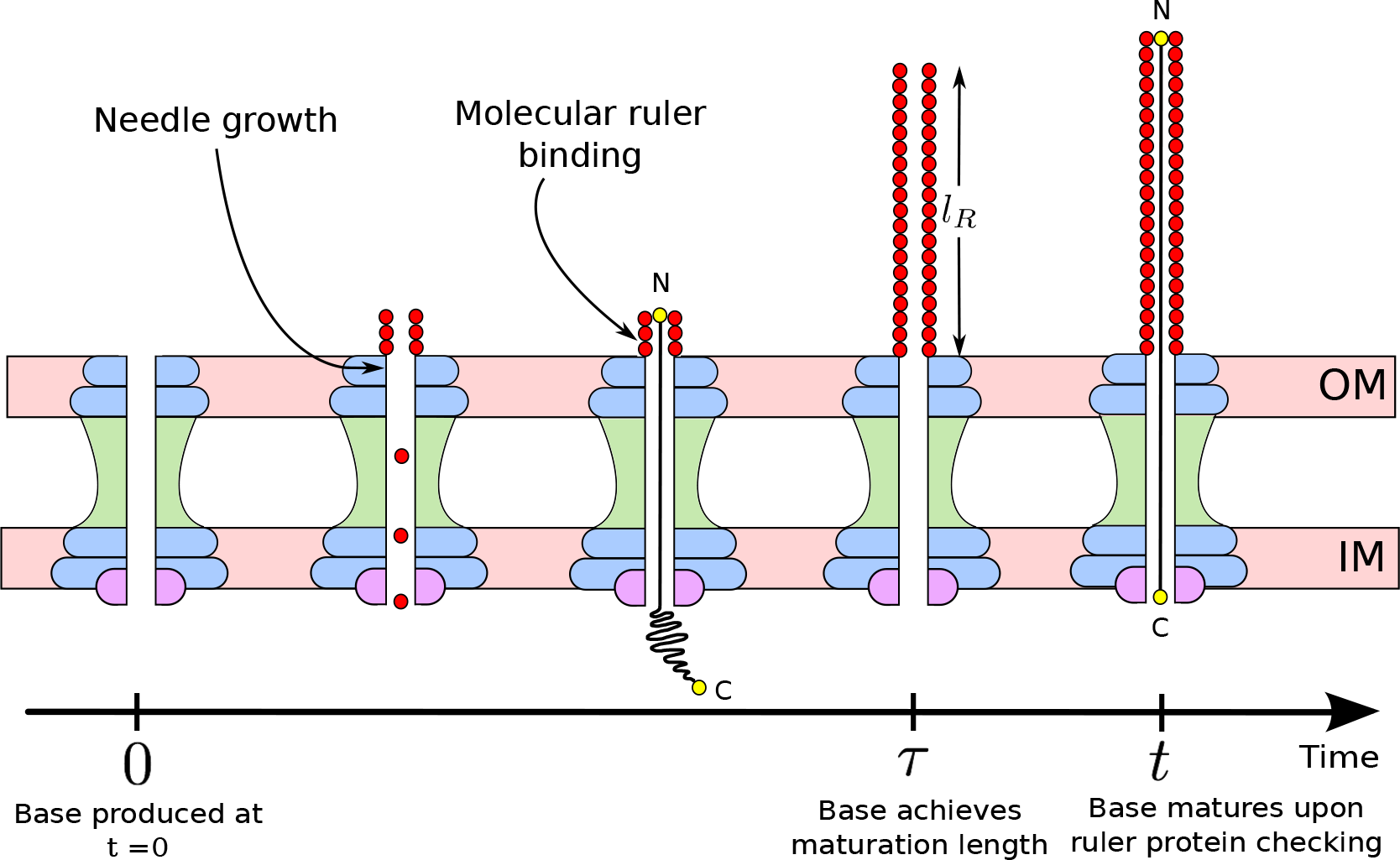
Figure 1 is a schematic of the growth of T3SS injectisome using an exact molecular ruler. A base which is produced at time *t* = 0, binds needle proteins and periodically checks for its needle length *L* with the effective ruler length *l*_*R*_. At time *τ* the needle achieves a length appropriate for maturation, *i.e. L* = *l*_*R*_, and the base matures at some later time *t* after exactly one ruler protein binding.

### Mathematical derivation

To develop a mathematical model for the ruler mechanism we considered the dynamics of the main constituent proteins of the system, namely the base, the needle (or the hook) proteins and the ruler protein. We represent the average numbers of the immature bases, needle proteins and ruler proteins as *B*, *O*, and *R* respectively. Since the flagellar hook as well as the injectisome needle, assemble outside the cell, we use the same notation, *O*, to denote both the hook protein and the needle protein. The assembly of these structures involves the following key processes: synthesis or production of the constituent proteins in the bacterial cell, export of the needle subunits and the ruler proteins by the base, and dilution of these proteins from cell division. In our derivation *Q*’s denote the production or synthesis rates whereas λ’s denote the dilution rates of the corresponding proteins. *β*_*O*_ and *β*_*R*_ are the rate at which the needle and the ruler proteins bind to the base complex and are exported, respectively. As seen in our previous work [13], we use a system of ordinary differential equations to represent the dynamics of the constituent proteins in the system (see the Supporting Information for more details).

We assume that the individual stochastic processes involved in the formation of the needle (namely the production, dilution and binding of the constituent proteins) are statistically independent, and we ignore any time delays that may arise due to synthesis or export of the proteins during the assembly process. We also assume that the system is already at steady state, *i.e.* the average values of *B*, *O* and *R* proteins are constant in time. Keeping these assumptions in mind, imagine a base being produced at *t* = 0. The probability that this base stops growing after achieving a certain needle length *L* can be written as:

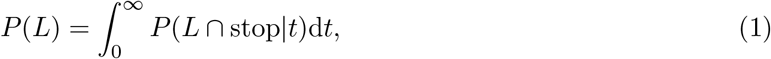

where *P*(*L*∩stop|*t*) is the probability density that the needle is of length *L* and the base stops growing it at time *t*. This can happen in two ways: either the base remains intact until time *t* and achieves maturation exactly at time *t* or the base stays immature until *t* and is lost due to dilution at time *t*. Mathematically, this can be written as:

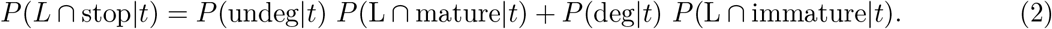

*P*(deg|*t*) represents the probability that the base is lost due to cell dilution at time *t*, whereas *P*(undeg|*t*) represents the probability that the base remains undiluted at time *t*. Both *P*(deg|*t*) and *P*(undeg|*t*) depend on a single rate parameter λ_*B*_. *P*(L ∩ mature|*t*) is a joint probability of the following three processes: the base achieves the maturation length *l*_*R*_ at time *τ* < *t*, there is one ruler protein binding event at time *t*, and the base continues to grow the needle until the ruler protein “checks” its length. *P*(L ∩ immature|*t*) involves dilution at time *t* and incorporates two scenarios: one where the length of the needle *L* is less than the required length for maturation at time *t* and a second where the base achieves the maturation length at time *τ* and no ruler checking events occur between *τ* and *t*. The detailed mathematical derivation can be found in the Supporting Information. The probability distribution for lengths in this model is:

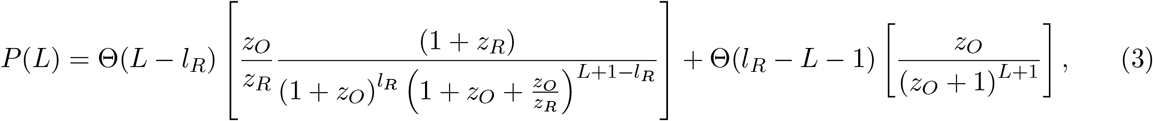

where *z*_*O*_ = λ_*B*_/*β*_*O*_*O* and *z*_*R*_ = λ_*B*_/*β*_*R*_*R*. The Heaviside theta functions ensure that the term associated with mature needles does not have lengths *L* < *l*_*R*_ and vice-versa.

For the bacteria to have most of its needles mature, it should be able to produce and bind enough needle proteins before it dilutes, which means *β*_*O*_*O* ≫ λ_*B*_ or *z*_*O*_ ≪ 1. Similarly, the rate at which the ruler protein checks the needles should also be sufficiently large or else dilution would dominate over maturation. At the same time it would be energetically inefficient for the bacteria to produce a large number of ruler proteins to check the needle too many times, so 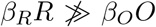. This leaves us with *z*_*R*_ ≲ *z*_*O*_. We used equation [3] and the ignored contributions from terms 𝒪(*z*^2^) or higher to obtain the average and variance in needle lengths (see the Supporting Information):

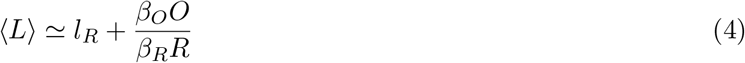

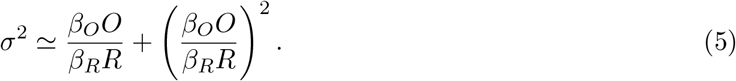

According to equations [4] and [5], the average length increases linearly with the length of the molecular ruler, whereas the variance is independent of the ruler protein length and only depends on the average number of ruler and outer proteins present in the bacterial cell.

### Comparison between model and stochastic simulation

In our derivation above, we made a number of simplifying assumptions (*e.g*. that the fluctuations in values of *B*, *O* and *R* are uncorrelated) that might not hold true in a bacterial cell. To test these assumptions, we performed stochastic simulations using the Doob-Gillespie algorithm [19]. In the simulations, the constituent proteins were treated as independent “agents” that interact with one another to form the needle (or the hook) [13, 20–23]. Each base has an arbitrary number of needle proteins associated with it. We keep track of the number of unbound constituent proteins, number of mature and immature bases, as well as the number of outer proteins attached to a given base. The precise values of the parameters used in this model have not been determined experimently, so for the purposes of comparison with our analytical results we chose parameter values subjected to reasonable constraints. Note that in our model all lengths are measured in terms of number of proteins. We used the structural details in [24] to convert the lengths to nanometers.

Figure 2 shows the comparison the mathematical model and the stochastic simulation at steady state in terms of experimentally relevant quantities such as average length, variance in length and ruler production rate. Note that in figure 2 we do not plot the solid lines for the entire range of parameters because the assumptions of our model are not valid in these regions (see Suppoting Information). In figure 2A, we observe a linear dependence of the average needle length on the ruler protein length for different ruler production rates, predicted as per equation (4). We notice that the average needle length increases in response to increase in the ruler length only upto a certain critical value 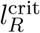, beyond which it remains constant. The constant length is a consequence of the fact that the bases go from being mostly mature, for instance, in figure 2A 100% of the bases are mature at *l*_*R*_ = 10 nm whereas only 40% of the bases are mature beyond 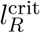 for *Q*_*R*_ = 5.0 molecules/sec. The average length in this case is close to ratio of production rates of the needle protein to that of the base (*Q*_*O*_/*Q*_*B*_) (in terms of number of proteins).

**Figure 2:**
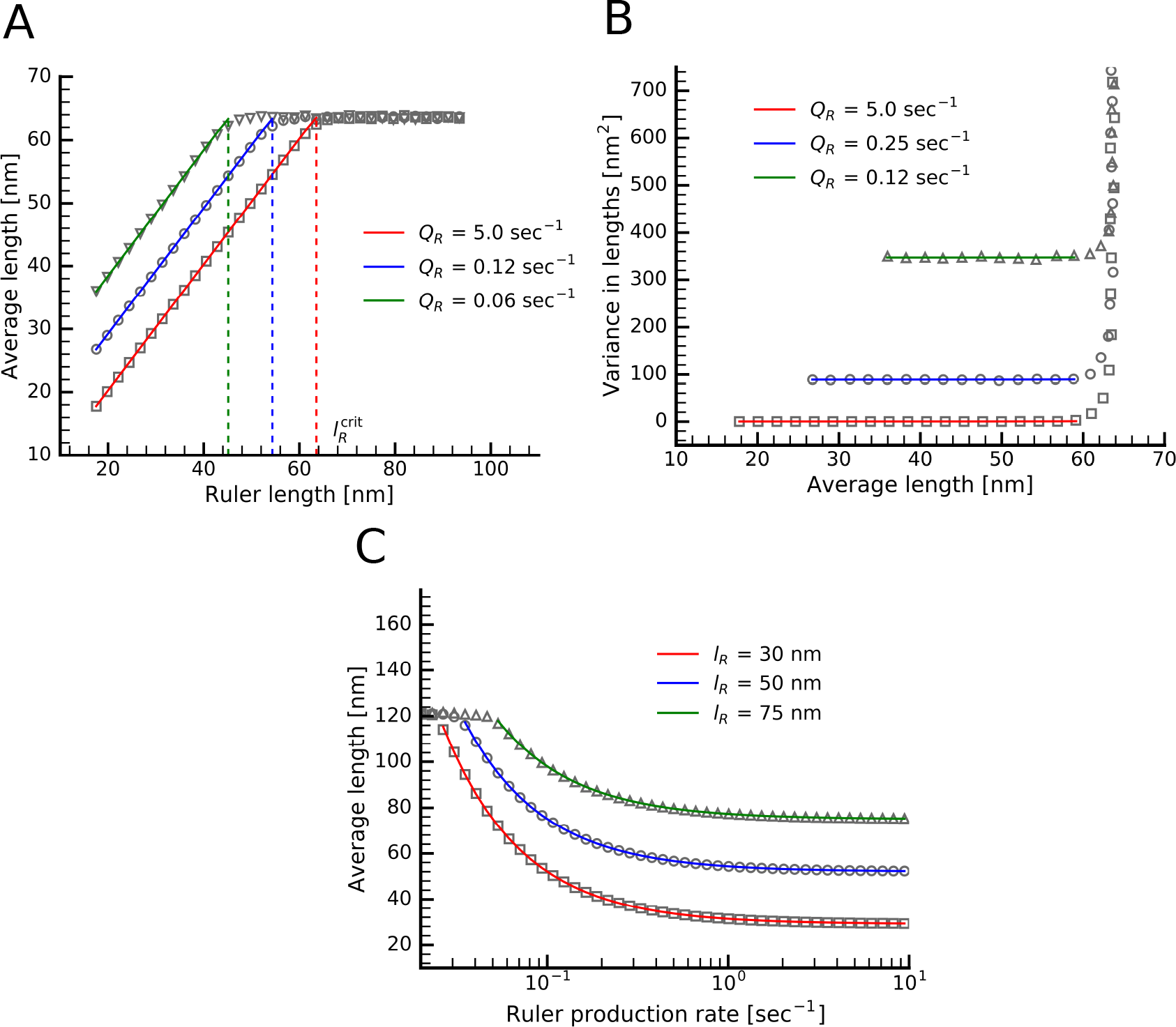
Figure compares the average and variance in lengths obtained from the mathematical model and the stochastic simulation. Figure 2A shows that average needle length increases linearly with the ruler length upto a certain value of ruler length 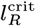, beyond which it remains constant and is approximately equal to *Q*_*O*_/*Q*_*B*_ (in terms of number of proteins). In Figure 2B we see that upon varying ruler length, the variance remains independent of the average needle length. In 2A and 2B the lines represent results from mathematical model for three different ruler production rates, red: *Q*_*R*_ = 5.0 sec^−1^, blue: *Q*_*R*_ = 0.12 sec^−1^ and green: *Q*_*R*_ = 0.06 sec^−1^, whereas the points represent the values from stochastic simulation for the corresponding ruler production rates. Figure 2C shows how the average length varies upon varying the ruler production rate. In figure 2C, the lines represent the results from mathmatical model for three different effective ruler lengths values, red: *l*_*R*_ = 30 nm, blue: *l*_*R*_ = 50 nm and green: *l*_*R*_ = 75 nm.

According to equation (5), the variance is independent of average length and depends only on the number of ruler and needle proteins. In figure 2B, we can see that upon varying *l*_*R*_, the variance remains constant with respect to the average length for variety of ruler production rates. Figure 2B also shows exceedingly large variances for extremely long ruler lengths. When the ruler proteins are extremely long, the percentage of immature bases increases, giving needles of varying lengths. The fact that the variance is independent of average length when most of the bases are mature makes the ruler model a unique mechanism. In [13] we found that the variance depends quadratically with the average length, and since the number of inner-rod proteins required to complete the inner-rod is more or less constant for any given species, there is no way to increase the length without entailing the quadratic increase in variance. However, in case of the (exact) ruler mechanism, the process of achieving the maturation length is completely deterministic. Thus the variance does not depend on the value of *l*_*R*_, and depends only on the rate parameters, *β*_*O*_*O* and *β*_*R*_*R*, making it possible to increase the average length by using longer rulers without changing the rate parameters of the ruler and needle proteins, leaving the variance unaltered (see equation 4 and 5). Thus the ruler mechanism could provide a much more tighly regulated needle length distribution than substrate switching.

Figure 2C shows the change in average length when the ruler protein production rate is varied. For large values of ruler protein concentration, the checking rate increases and hence the needles achieve maturation at a length close to the ruler length. Decreasing the ruler protein production reduces the frequency of ruler checking, thus allowing the needles to grow beyond the ruler length before they become mature. At very low production rates of the ruler protein, there are so few rulers compared to the number of bases that most bases remain immature and the average needle length in the population no longer depends on *l*_*R*_.

### Experimental data

In [10] the authors changed the ruler protein length by adding residues to the wild type ruler protein YscP in *Y. pestis* and saw a linear relation between the ruler length and the average needle lengths. A similar study was conducted by examining the flagellar hook lengths when the length of the FliK was varied by adding chimeric residues in [11]. More recently, researchers have identified InvJ (in *S. enterica*), a portein homologuous to YscP as well as FliK, and studied the effect of varying its length on the average length of T3SS needles [12]. The key result of these experiments was that the average length increased linearly as the length of the molecular ruler, giving strong evidence for the ruler mechanism. We obtained the variance the needle and hook lengths from the data in these studies. Figures 3A and 3B show that variance is not independent of the average length, but rather increases linearly with the average length. Our regression analysis confirms that there was no statistical significance for a quadratic relationship between these two quantities (see the Supporting Information). Although a linear relationship between variance and needle length allows a tighter control over needle lengths as compared to quadratic relationship, this experimental data is not consistent with the exact ruler model.

**Figure 3:**
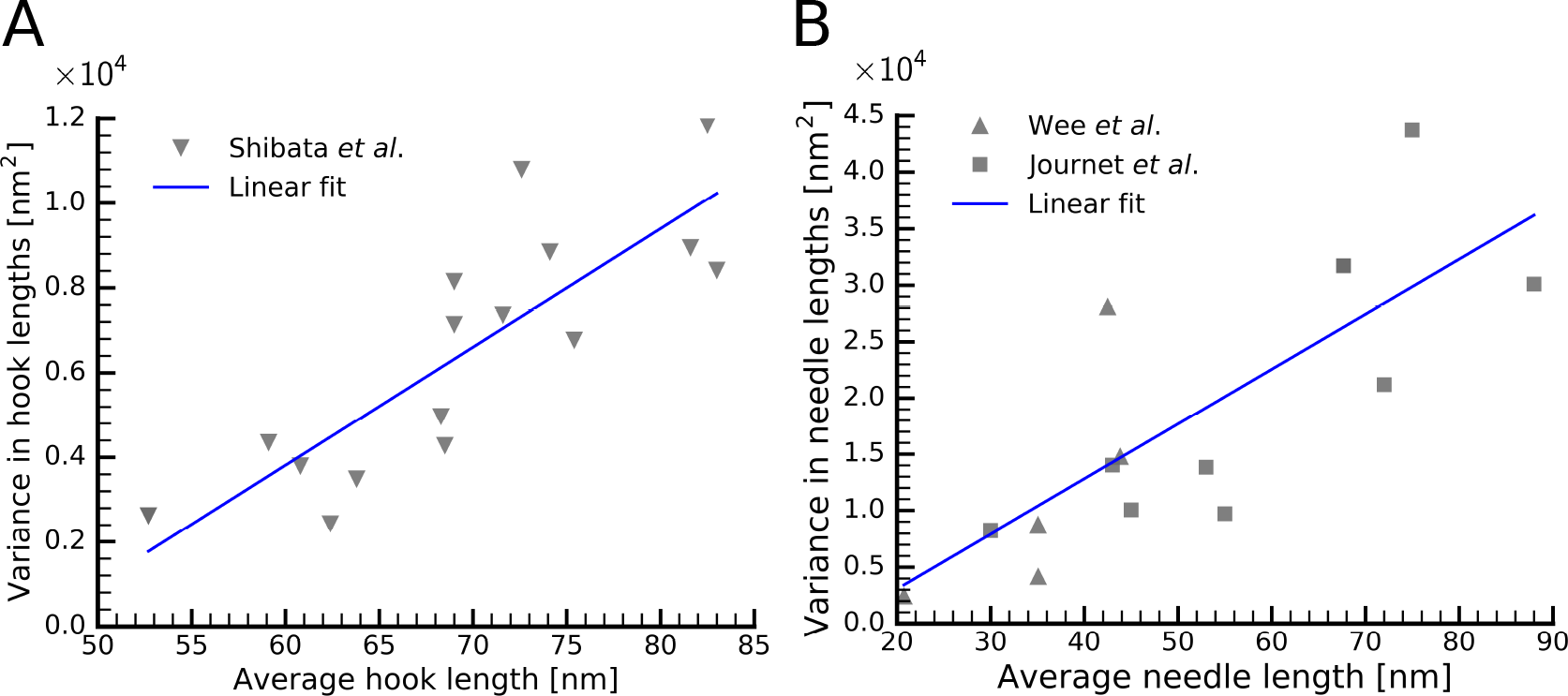
Figure 3 shows the dependence of variance on the average length in flagellar hooks in [11] and in T3SS injectisome in [12] and [10]. Our regression analysis shows no statistical significance for a quadratic relation between the variance and the average length, the points denote the experimental data whereas the lines represent the linear fit.

### Ruler length with an error-prone ruler

One of the assumptions in our initial model was that the interaction of the C-terminus of the ruler protein with the gate protein occurs at a fixed length. In reality, this interaction depends on conformational changes that arise as a consequence of the folding of the ruler protein, resulting in uncertainties in the measurement process. To incorporate this effect in our simulation, we allowed the length of the ruler involved in every binding interaction to fluctuate in such a way that maturation probability obeys a logistic distibution (see Supporting Information) [15]. This variability in the effective ruler length makes it a possible for the base achieve maturation for a needle length that might have been too short for maturation in the exact ruler model. This effect is not straightforward to incorporate in the analytical calculation for the probability distribution of needle lengths, and thus in the analysis that follows we focus on simulation results.

Figure 4 shows the average and variance in lengths for a logistic ruler with a fixed error in length. In figure 4A we observe that varying the length of the error-prone ruler (keeping all other parameters fixed) leads to a linear relationship between the average length and the ruler length, which is what observed in case of the exact ruler as well. In figure 4B we can see the change in average lengths when the ruler production rate is varied. For lower production rates, only a few rulers are being produced in comparison to the number of bases available, giving us the same average lengths as obtained using the exact ruler. But when we increase the ruler production rates, the average lengths in case of a logistic ruler decrease more rapidly than they did in case of an exact ruler. This is because, at higher production rates, even though maturation at small values of *L* is less likely to occur, there are enough ruler checking events to ensure maturation even for needles with *L* significantly less than *l*_*R*_. Note that this effect is prominent even at moderate values of *Q*_*R*_. In figure 4C we observe that varying the length of the error-prone ruler (same conditions as in Figure 4A) leads to a linear relationship between the variance and the average length. A logistic ruler allows for maturation events at much smaller neeedle lengths, giving a large number of needles whose lengths are less than *l*_*R*_. This skews the distribution of needle lengths resulting in larger variances. Note that variance obtained through the simulation is smaller than that observed in the experiments, which could be due a number of experimental uncertainties that were not incorporated in our simulations.

**Figure 4:**
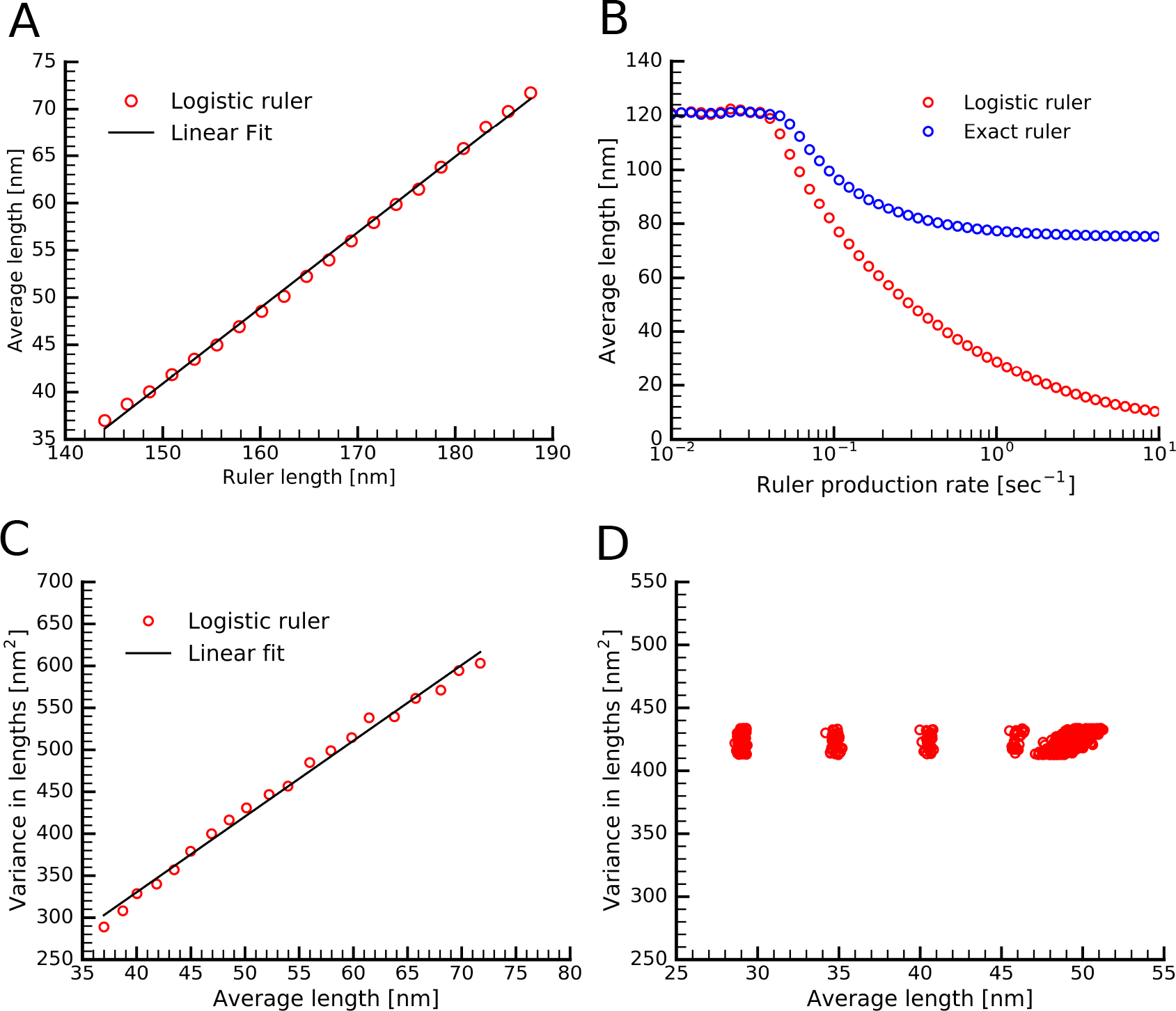
Figure 4 shows average and variance in lengths using a logistic or “error-prone” ruler. Figure 4A shows that the average length increase linearly with the ruler length. Figure 4B shows that the average length of the needle is approximately equal in case of exact and the logistic ruler for relatively lower ruler production values. However, for larger ruler production values, even though the probability of having a short length an effective ruler, the chances of a needle with such a length achieving maturation increases dramatically. In 4C we see that variance in length increases linearly with the average length when the ruler length is increased. Since the logistic ruler allows maturation relatively smaller needles, this skews the needle length distribution, increasing the varaince in lengths. The ruler lengths were varied from 120–200 nm with a fixed error of 30 nm in length, *Q*_*R*_ = 7.5 and *Q*_*O*_ = 3.75. Figure 4C shows that as per a logistic ruler it is possible to vary the ruler length without increasing the variance. *l*_*R*_ = 65–100 nm with a relative error in length of 10%, *Q*_*R*_ = 0.25−2.75 and *Q*_*O*_ = 1.0−3.25. Figure 4D variance remains constant over a range of ruler lengths for different *Q*_*R*_ and *Q*_*O*_ values.

As discussed earlier, for an exact ruler the variance in length is independent of the ruler length itself, and depends only on the needle and ruler protein concentrations and export rates. We should note that Figures 4A and 4C represent a typical experimental situation in which the ruler protein length is changed *via* deletion of amino acid residues in the protein, or insertion of new residues into a specific region [10–12]. In this scenario, we find that the variance increases linearly with the average needle length, which is also what is observed experimentally (Figures 3 and 4C). During the course of evolution, however, it would be possible for both the length of the ruler protein and the synthesis rates of the ruler and needle proteins to vary simultaneously. We thus tested whether it would be possible to find parameter sets where the increase in variance seen in Fig. 4C could be offset by varying other parameters. Interestingly, we found that it is indeed possible to increase the average length of the needles *without* significantly changing the variance (Fig. 4D). While more complex than the case with an exact ruler, it is thus nonetheless possible for an error-prone ruler to exhibit an independence of the average and variance in the length distribution.

## Discussion

The T3SS and flagellar hook are large and complex nanomachines that are crucial to bacterial cell function, pathogenesis and adaptation to changing environments. One key aspect of these homologous structures is the large extracellular channel that must be constructed for both of them. The lengths of these channels have to be extremely precise; if they are too short or too long, that may negatively influence the corresponding fucntion. For instance, in the case of T3SS needles, if the needles are too short then they would be unable to get past the lipopolysccharide (LPS) layer present in the extracelluar region of the bacterium, thus failing to inject the effector proteins into the host cell. On the other hand if the needles are too long, then they will likely break from shear stress or lead to energetically inefficent transport of proteins through an excessively long channel. Similarly, a flagellar hook of an inappropriate size might lead to poor mechanical properties for locomotion.

The substrate switching and ruler mechanisms are the two most popular and well-studied explanations for length control in these structures. The substrate switching mechanism requires an inner-rod present inside the base to be completed in order to form a mature needle, whereas according to the ruler mechanism, the bacteria “measures” the length of these structures with a dedicated ruler protein. While there is some experimental evidence for these mechanisms, until recently several important questions remained unanswered. For instance, is there a quantitative consitency of the proposed mechanism with experiments? What are the advantages, from an evolutionary perspective, to adapt different length control mechanisms? What future experiments could be done to further investigate these mechanisms?

In our previous work, we constructed a mathematical model for length control as per the substrate switching mechanism and compared our predictions with the available experimental data for *Salmonella*. Our model was able to explain the length distibution in the needle lengths obtained in [8], and we were also able to predict the number the of PrgJ proteins required to complete the inner-rod and found that our prediction was consistent with data from mass spectrometry [14]. In the current work we developed a mathematical model for the ruler mechanism. Interestingly, we found that an exact ruler, which is perhaps most commonly envisioned for length measurement according to the ruler mechanism, does not explain the experimental data, and that a logistic ruler, which has a measurement uncertainty associated with it, is consistent with the available experimental data [10–12]. Interestingly, an error-prone ruler allows bacterial cells to regulate the variance to a desired value by adapting the ruler length and ruler and needle proteins synthesis rates simultaneously. This allows the ruler mechanism to exert a greater degree of control over the length of the structure than the substrate switching mechanism, particularly when the structures themselves are long. This higher degree of control, however, comes at a fairly high energetic cost. In particular, once ruler proteins are secreted, they are lost, and obtaining a low variance requires many ruler protein “measurements” for every base that becomes mature. In contrast, the substrate switching model involves a constant number of inner-rod proteins for each base, and if this number is low (around six or so, as predicted by our model and observed in experimental data [13, 14]), then the energetic cost of control in this case is relatively low. We thus might expect the substrate switching mechanism to be favored as a low-energy control strategy for short structures, whereas the ruler protein mechanism might be favored for longer structures where the lower cost of the substrate switching mechanism would be outweighed by the massive increase in variance that it entails as the needles or hooks become longer.

In addition to providing insight into the evolutionary trade-offs of various control strategies, our quantitative models suggest several experiments that could provide further insight into the control mechanisms themselves. For instance, researchers have studied the effect of varying the length of ruler proteins such as YscP (in *Yersinia*) and FliK (in *Salmonella*) on the length of needles and flagellar hooks respectively, so an experiment where one changes the concentration of the ruler proteins by means of titrable promotors for the wild type as well as for ruler length variants of these ruler proteins would help find evidence for the ruler mechanism (See figure 2C and 4C). A similar experiment where one changes the concentration of the inner-rod protein PrgJ in *Salmonella* can provide evidence for the substrate switching mechanism [13]. In addition to the inner-rod protein PrgJ, researchers have also identified a homologue of ruler protein, InvJ, in *Salmonella* [12]. By varying the concentration of InvJ as well as PrgJ, one can find whether the substrate switching or the ruler mechanism is operational for that species. It is clear that a combination of quantitative models and experiments will be crucial for building a complete understanding of the assembly and regulation of these massive extracellular structures.

