## Supporting Information for "Robustness and the evolution of length control strategies in the type III secretion system and flagellar hook"

### Probability distribution for needle lengths

As per the ruler model, the ruler protein of fixed length $l_R$ periodically binds to the base allowing it to "compare" its needle length to that of the ruler protein. If the needle length is greater than or equal to that of the ruler protein then the needle stops growing and base achieves maturation. Imagine a base that is born at time $t = 0$. The probability that this base has a needle of length $L$ at the steady state is:

$$P(L) = \int_0^\infty P(L \cap \text{stop}|t)dt, \tag{1}$$

where $P(L \cap \text{stop}|t)$ is the probability that the base is of length $L$ and it stops growing the needle at time $t$. This can happen if the base remains intact until time $t$ and achieves maturation exactly at time $t$ or that the base stays immature until and $t$ and is lost due to cell division at time $t$. Mathematically, this could be written as,

$$P(L \cap \text{stop}|t) = P(\text{undeg}|t)\ P(L \cap \text{mature}|t) + P(\text{deg}|t)\ P(L \cap \text{immature}|t), \tag{2}$$

$P(\text{undeg}|t)$ and $P(\text{deg}|t)$ are the same expressions as we had in case of substrate switching [1],

$$P(\text{undeg}|t) = \mathrm{e}^{-\lambda_B t}$$
$$P(\text{deg}|t) = \lambda_B\ \mathrm{e}^{-\lambda_B t}.$$

where $\lambda_B$ is the degradation rate of the base due to cell dilution.

The process where a base has a needle of length $L$ and is mature at time $t$ can can only occur if the base achieves the maturation length at an intermediate time $\tau$ and there is exactly one ruler protein binding event between $\tau$ and $t$;

$$P(L \cap \text{mature}|t) = \int_0^t \underbrace{\left(\mathrm{e}^{-\beta_O O \tau} \frac{(\beta_O O)^{l_R} \tau^{l_R - 1}}{(l_R - 1)!}\right)}_{l_R \text{ needle protein exports in time } \tau} \underbrace{\beta_R R \mathrm{e}^{-\beta_R R(t-\tau)}}_{1 \text{ ruler protein at } t}$$

$$\underbrace{\left(\mathrm{e}^{-\beta_O O(t-\tau)}\frac{\left[\beta_O O(t-\tau)\right]^{L-l_R}}{(L-l_R)!}\right)}_{L-l_R \text{ needle protein exports between } \tau \text{ and } t} \quad d\tau \tag{3}$$

where $\beta_O, \beta_R$ are the binding rate of the outer and ruler proteins respectively, and $O, R$ are the steady state values of the concentration of the outer and ruler proteins respectively. In equation (3), the first term in the integral denotes the probability that the base achieves the length $l_R$ at time $\tau$, the second term denotes the probability that there is one checking event at time $t$ such that $t > \tau$ , and the third term incorporates the fact that the base could have $(L - l_R)$ needle protein binding events after achieving the maturation length at $\tau$ and before the next ruler protein binding at $t$, at which point the base becomes mature.

A base can remain immature if (1) it has less than $l_R$ needle proteins bound to it at time $t$, the time at which it degraded, *or* (2) it has achieved the maturation length at some intermediate time $\tau$ and there are no ruler protein bindings between the times $\tau$ and $t$.

$$P(L \cap \text{immature}|t) = \underbrace{\mathrm{e}^{-\beta_O O t}\frac{(\beta_O O t)^L}{L!}}_{L \text{ needle protein exports in time } t} + \int_0^t \underbrace{\left(\mathrm{e}^{-\beta_O O\tau}\frac{(\beta_O O)^{l_R}\tau^{l_R-1}}{(l_R-1)!}\right)}_{l_R \text{ needle protein exports in } t}$$
$$\underbrace{\mathrm{e}^{-\beta_R R(t-\tau)}}_{0 \text{ ruler protein exports between } \tau \text{ and } t} \underbrace{\left(\mathrm{e}^{-\beta_O O(t-\tau)}\frac{\left[\beta_O O(t-\tau)\right]^{L-l_R}}{(L-l_R)!}\right)}_{L-l_R \text{ needle protein exports between } \tau \text{ and } t} \quad d\tau \tag{4}$$

In equation (4), scenario (1) is represented by the term outside the integral denotes the probability that there are $L$ needle protein export events until time $t$. Scenarios is represented by the other two terms in the equation, the first term inside the integral denotes the probability that there are $l_R$ needle protein binding events until time $\tau$, the second terms the probability that there are no ruler checking events between $\tau$ and $t$, and the third term denotes the probability that there are $L - l_R$ needle protein export events between $\tau$ and $t$.

Combining equations (1)–(4),

$$P(L) = \Theta(L-l_R)\int_0^\infty dt\ \mathrm{e}^{-\lambda_B t}\int_0^t d\tau \left(\mathrm{e}^{-\beta_O O\tau}\frac{(\beta_O O)^{l_R}\tau^{l_R-1}}{(l_R-1)!}\right)\beta_R R\mathrm{e}^{-\beta_R R(t-\tau)}\left(\mathrm{e}^{-\beta_O O(t-\tau)}\right.$$
$$\left.\frac{\left[\beta_O O(t-\tau)\right]^{L-l_R}}{(L-l_R)!}\right) + \Theta(l_R-L-1)\int_0^t dt\ \lambda_B\ \mathrm{e}^{-\lambda_B t}\mathrm{e}^{-\beta_O O t}\frac{(\beta_O O t)^L}{L!}$$
$$+\Theta(L-l_R)\int_0^\infty dt\ \lambda_B \mathrm{e}^{-\lambda_B t}\int_0^t d\tau\ \left(\mathrm{e}^{-\beta_O O\tau}\frac{(\beta_O O)^{l_R}\tau^{l_R-1}}{(l_R-1)!}\right)\mathrm{e}^{-\beta_R R(t-\tau)}$$

$$\left( \mathrm{e}^{-\beta_O O(t-\tau)} \frac{\left[\beta_O O(t-\tau)\right]^{L-l_R}}{(L-l_R)!} \right) \tag{5}$$

The Heaviside theta functions, $\Theta(L - l_R)$ and $\Theta(l_R - L - 1)$, ensure that the term associated with mature needles does not have lengths $L < l_R$ and vice-versa. We performed the integration in the above expressions using Gamma functions and verified our calculations using Mathematica (v11.0.0.0),

$$P(L) = \Theta(L - l_R) \left[ \frac{(\beta_O O)^L (\lambda_B + \beta_R R)}{(\lambda_B + \beta_O O)^{l_R} (\lambda_B + \beta_O O + \beta_R R)^{L+1-l_R}} \right] + \Theta(l_R - L - 1) \left[ \frac{\lambda_B (\beta_O O)^L}{(\lambda_B + \beta_O O)^{L+1}} \right] \tag{6}$$

Rearranging the terms in equation (6), we get:

$$\begin{aligned} P(L) &= \frac{(\lambda_B + \beta_R R)(\lambda_B + \beta_O O + \beta_R R)^{l_R - 1}}{(\lambda_B + \beta_O O)^{l_R}} \left( \frac{\beta_O O}{\lambda_B + \beta_O O + \beta_R R} \right)^L \Theta(L - l_R) \\ &+ \frac{\lambda_B}{\lambda_B + \beta_O O} \left( \frac{\beta_O O}{\lambda_B + \beta_O O} \right)^L \Theta(l_R - L - 1) \\ &= \left( \frac{\lambda_B + \beta_R R}{\lambda_B + \beta_O O + \beta_R R} \right) \left( \frac{x_2}{x_1} \right)^{l_R} x_1^L \, \Theta(L - l_R) + \frac{\lambda_B}{\lambda_B + \beta_O O} x_2^L \, \Theta(l_R - L - 1) \\ &= (1 - x_1) \left( \frac{x_2}{x_1} \right)^{l_R} \Theta(L - l_R) \, x_1^L + (1 - x_2) \, \Theta(l_R - L - 1) \, x_2^L \end{aligned} \tag{7}$$

where

$$\begin{aligned} x_1 &= \frac{\beta_O O}{\lambda_B + \beta_O O + \beta_R R} = \frac{1}{1 + (\lambda_B/\beta_O O) + (\beta_R R/\beta_O O)} = \frac{1}{1 + z_O + z_O/z_R} \\ x_2 &= \frac{\beta_O O}{\lambda_B + \beta_O O} = \frac{1}{1 + (\lambda_B/\beta_O O)} = \frac{1}{1 + z_O} \end{aligned}$$

with

$$z_O = \lambda_B/\beta_O O \,, \qquad z_R = \lambda_B/\beta_R R \,, \qquad z_O/z_R = \beta_R R/\beta_O O$$

In equation (7), length distribution $P(L)$ can be expressed in terms of $(x_1, x_2)$ or $(z_O, z_R)$ for a given $l_R$.

### Normalization of $P(L)$

$$\begin{aligned} \sum_{L=0}^{\infty} P(L) &= (1 - x_1) \left( \frac{x_2}{x_1} \right)^{l_R} \sum_{L=l_R}^{\infty} x_1^L + (1 - x_2) \sum_{L=0}^{l_R - 1} x_2^L \\ &= (1 - x_1) x_2^{l_R} \frac{1}{1 - x_1} + (1 - x_2) \frac{1 - x_2^{l_R}}{1 - x_2} = 1 \end{aligned}$$

where

$$\sum_{L=l_R}^{\infty} x_1^L = \frac{x_1^{l_R}}{1 - x_1} \qquad \text{and} \qquad \sum_{L=0}^{l_R - 1} x_2^L = \frac{1 - x_2^{l_R}}{1 - x_2}$$

### Calculation of average length and variance in lengths

We can obtain the average and the variance by calculating the 1st and the 2nd moments of the probability distribution $P(L)$.

$$
\begin{aligned}
\langle L \rangle &= \sum_{L=0}^{\infty} P(L)L \;=\; (1-x_1)\left(\frac{x_2}{x_1}\right)^{l_R} \sum_{L=l_R}^{\infty} L x_1^L + (1-x_2)\sum_{L=0}^{l_R-1} L x_2^L \\
&= (1-x_1)\left(\frac{x_2}{x_1}\right)^{l_R} x_1 \frac{d}{dx_1}\left(\sum_{L=l_R}^{\infty} x_1^L\right) + (1-x_2)x_2 \frac{d}{dx_2}\left(\sum_{L=0}^{l_R-1} x_2^L\right) \\
&= (1-x_1)\left(\frac{x_2}{x_1}\right)^{l_R} x_1 \frac{d}{dx_1}\left(\frac{x_1^{l_R}}{1-x_1}\right) + (1-x_2)x_2 \frac{d}{dx_2}\left(\frac{1-x_2^{l_R}}{1-x_2}\right) \\
&= x_2^{l_R}\left(l_R + \frac{x_1}{1-x_1}\right) - l_R x_2^{l_R} + x_2 \frac{1-x_2^{l_R}}{1-x_2} \\
&= \frac{x_2}{1-x_2} - x_2^{l_R}\left(\frac{x_2}{1-x_2} - \frac{x_1}{1-x_1}\right) \\
&= \frac{1}{z_O}\left(1 - \frac{x_2^{l_R}}{1+z_R}\right) \;=\; \frac{1}{z_O}\left[1 - \frac{1}{(1+z_O)^{l_R}(1+z_R)}\right]
\end{aligned} \tag{8}
$$

where

$$
\frac{x_1}{1-x_1} = \frac{z_R}{z_O(z_R+1)} \qquad \text{and} \qquad \frac{x_2}{1-x_2} = \frac{1}{z_O}
$$

and

$$
\left\langle L^2 \right\rangle = \sum_{L=0}^{\infty} P(L)L^2 = \sum_{L=0}^{\infty} P(L)L(L-1) + \sum_{L=0}^{\infty} P(L)L = \sum_{L=0}^{\infty} P(L)L(L-1) + \langle L \rangle
$$

where

$$
\begin{aligned}
\sum_{L=0}^{\infty} P(L)L(L-1) &= (1-x_1)\left(\frac{x_2}{x_1}\right)^{l_R} \sum_{L=l_R}^{\infty} L(L-1)x_1^L + (1-x_2)\sum_{L=0}^{l_R-1} L(L-1)x_2^L \\
&= (1-x_1)\left(\frac{x_2}{x_1}\right)^{l_R} x_1^2 \frac{d^2}{dx_1^2}\left(\sum_{L=l_R}^{\infty} x_1^L\right) + (1-x_2)x_2^2 \frac{d^2}{dx_2^2}\left(\sum_{L=0}^{l_R-1} x_2^L\right) \\
&= (1-x_1)\left(\frac{x_2}{x_1}\right)^{l_R} x_1^2 \frac{d^2}{dx_1^2}\left(\frac{x_1^{l_R}}{1-x_1}\right) + (1-x_2)x_2^2 \frac{d^2}{dx_2^2}\left(\frac{1-x_2^{l_R}}{1-x_2}\right) \\
&= x_2^{l_R}\left[l_R(l_R-1) + \frac{2l_R x_1}{1-x_1} + \frac{2x_1^2}{(1-x_1)^2}\right] + \frac{2x_2^2}{(1-x_2)^2} - x_2^{l_R}\left[l_R(l_R-1) + \frac{2l_R x_2}{1-x_2} + \frac{2x_2^2}{(1-x_2)^2}\right] \\
&= \frac{2x_2^2}{(1-x_2)^2} + 2x_2^{l_R}\left[l_R\left(\frac{x_1}{1-x_1} - \frac{x_2}{1-x_2}\right) + \frac{x_1^2}{(1-x_1)^2} - \frac{x_2^2}{(1-x_2)^2}\right] \\
&= \frac{2}{z_O^2} + 2x_2^{l_R}\left[-\frac{l_R}{z_O(1+z_R)} + \frac{z_R^2}{z_O^2(1+z_R)^2} - \frac{1}{z_O^2}\right] \\
&= \frac{2}{z_O^2}\left[1 - \frac{1}{(1+z_O)^{l_R}(1+z_R)}\left(2 + l_R z_O - \frac{1}{1+z_R}\right)\right]
\end{aligned}
$$

Since from $\langle L \rangle$ in Eq. (8), we have

$$\frac{1}{1+z_R} = (1+z_O)^{l_R}(1 - z_O \langle L \rangle)$$

$$\begin{aligned}
\sum_{L=0}^{\infty} P(L)L(L-1) &= \frac{2}{z_O^2}\left\{1 - \frac{(1+z_O)^{l_R}(1-z_O\langle L\rangle)}{(1+z_O)^{l_R}}\left[2 + l_R z_O - (1+z_O)^{l_R}(1-z_O\langle L\rangle)\right]\right\} \\
&= \frac{2}{z_O^2}\left[1 - (2 + l_R z_O)(1 - z_O\langle L\rangle) + (1+z_O)^{l_R}(1-z_O\langle L\rangle)^2\right]
\end{aligned}$$

Thus

$$\begin{aligned}
\sigma^2 &= \left\langle L^2 \right\rangle - \langle L\rangle^2 \\
&= \frac{2}{z_O^2}\left[1 - (2 + l_R z_O)(1 - z_O\langle L\rangle) + (1+z_O)^{l_R}(1-z_O\langle L\rangle)^2\right] + \langle L\rangle - \langle L\rangle^2 \\
&= \frac{2}{z_O^2}\left[(1+z_O)^{l_R} - (1 + l_R z_O)\right] \\
&\quad - \langle L\rangle\left\{\frac{4}{z_O}\left[(1+z_O)^{l_R} - 1\right] - (2l_R + 1)\right\} + \langle L\rangle^2\left[2(1+z_O)^{l_R} - 1\right] \\
&= a\langle L\rangle^2 - 2b\langle L\rangle + c \;=\; a\left(\langle L\rangle - \frac{b}{a}\right)^2 + \left(c - \frac{b^2}{a}\right) \qquad (9)
\end{aligned}$$

where $a$, $b$ and $c$ are

$$\begin{aligned}
a &= 2(1+z_O)^{l_R} - 1 \\
b &= \frac{2}{z_O}\left[(1+z_O)^{l_R} - (1 + \frac{l_R z_O}{2})\right] - \frac{1}{2} \\
c &= \frac{2}{z_O^2}\left[(1+z_O)^{l_R} - (1 + l_R z_O)\right] \\
\frac{b}{a} &= \frac{1}{z_O} - \frac{2 + (2l_R - 1)z_O}{2z_O[2(1+z_O)^{l_R} - 1]}
\end{aligned}$$

### Approximation in biologically relevant parameter regime

For the bacteria to have almost all needles mature, it should be able to produce and bind enough needle proteins, which means $\beta_O O \gg \lambda_B$ or $z_O \ll 1$. Furthermore, the needles should also be checked by attaching the ruler protein enough number of times or else it result in devastatingly large needles, which means $\beta_R R > \lambda_B$. At the same time it would be energetically inefficient for the bacteria to check the needle too many times, so $\beta_R R \not\gg \beta_O O$. This leaves us with $z_R \simeq z_O$. So to obtain the expressions for average needle length and variance in the parameter space where most needles are mature we can ignore terms $\mathcal{O}(z^2)$ or higher.

From equation (8) we have

$$\langle L \rangle = \frac{1}{z_O}\left[1 - \frac{1}{(1+z_O)^{l_R}(1+z_R)}\right]$$

Ignoring terms $\mathcal{O}(z^2)$ and higher

$$\begin{aligned}
&\simeq \frac{1}{z_O}\left[\frac{(1+l_R z_O)(1+z_R)-1}{(1+l_R z_O)(1+z_R)}\right] \\
&= \frac{1}{z_O}\left[\frac{l_R z_O + z_R}{1+l_R z_O + z_R}\right] \\
&\simeq l_R + \frac{z_R}{z_O} = l_R + \frac{\beta_O O}{\beta_R R}
\end{aligned} \tag{10}$$

Similarly we can obtain the variance by using the approximate values of $a, b, c$:

$$\begin{aligned}
a &= 2(1+z_O)^{l_R} - 1 & &\overset{z_O \ll 1}{=} 1 \\
b &= \frac{2}{z_O}\left[(1+z_O)^{l_R} - (1+\frac{l_R z_O}{2})\right] - \frac{1}{2} & &\overset{z_O \ll 1}{=} l_R - \frac{1}{2} \\
c &= \frac{2}{z_O^2}\left[(1+z_O)^{l_R} - (1+l_R z_O)\right] & &\overset{z_O \ll 1}{=} l_R(l_R - 1)
\end{aligned}$$

Using the approximate values of $a$, $b$, $c$, and $\langle L\rangle$ in equation (9), we get

$$\sigma^2 \simeq \frac{z_R}{z_O} + \left(\frac{z_R}{z_O}\right)^2 = \frac{\beta_O O}{\beta_R R} + \left(\frac{\beta_O O}{\beta_R R}\right)^2 \tag{11}$$

### Ordinary differential equations

In order to calculate the average the variance, we need to obtain the steady state values of the number base, the ruler and the needle proteins ($B$, $R$, and $O$ respectively), which are evaluated by solving a system of ordinary differential equations (ODE). To construct the ODE's, consider a base produced at time $t = 0$. Let $\tau$ denote the average time that the base requires to achieve the maturation length. The binding rate of the ruler protein is $\beta_R$, which means the average time lapsed between two consecutive ruler protein binding is $1/(\beta_R R)$. Thus, the average time required for maturation is:

$$\langle T\rangle = \tau + \frac{1}{\beta_R R}$$

If we assume that the ruler protein binding and needle protein binding processes to be completely independent of one another then $\tau$ is :

$$\tau = \frac{l_R}{\beta_O O}$$

giving,

$$\langle T\rangle = \frac{l_R \beta_R R + \beta_O O}{\beta_O \beta_R O R}. \tag{12}$$

This means that on average, after every $\langle T\rangle$ seconds a base achieves maturation and is lost from the pool of immature bases. Thus the ODE's for this system are:

$$\frac{\mathrm{d}B}{\mathrm{d}t} = Q_B - \lambda_B B - \frac{B}{\langle T\rangle} \tag{13}$$

$$\frac{\mathrm{d}R}{\mathrm{d}t} = Q_R - \lambda_R R - \beta_R R B \tag{14}$$

$$\frac{\mathrm{d}O}{\mathrm{d}t} = Q_O - \lambda_O O - \beta_O O B \tag{15}$$

The solution to these ODE's gives the steady state values of $O, R$ and $B$, thus we can obtain the average and the variance calculated in the previous section from equations (10) and (11).

### Stochastic simulations

We treated the bases as independent "agents" in our simulations. Each base has a number of needle proteins associated with it that can take any non-negative integer value. We maintained two separate populations of bases; immature bases, represented by $B$, and mature bases represented by $B'$ (total number of bases $B_{\text{tot}} = B + B'$). A base achieves maturation upon the first ruler protein checking event that occurs after its needle grows to a length that is greater than or equal to $l_R$. In addition to the number of bases we use two integers, $R$ and $O$, to keep track of the number of free ruler and needle proteins.

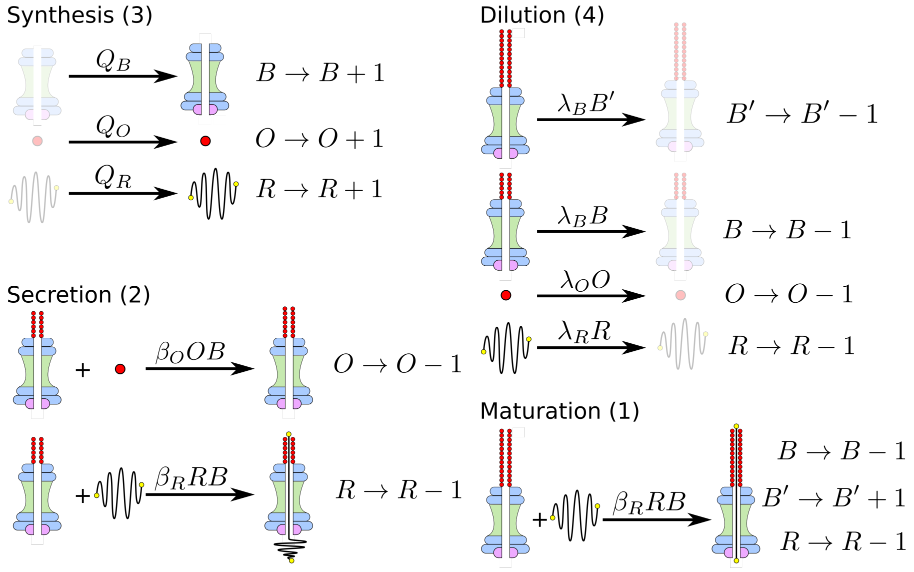


Figure 1: Schematic representation of the processes included in the simulation. Top left shows the synthesis of a base, a ruler protein, and a needle protein. Bottom left shows the ruler protein and needle needle protein. Top right shows the dilution of the constituent proteins from cell division, and bottom right shows maturation of a base. The propensities of these reactions are mentioned on the the arrows, whereas the equation on the right represents the change in the corresponding numbers when a particular reaction is chosen

Based on the assumption of the model, the possible chemical reactions that can occur between the bases, ruler, and needle proteins and their effect on the quantities described above are show in figure

1. We initialized our simulations with a set of immature bases with no needle proteins bound. When a base was synthesized, an immature base without any needle protein bound to it was added to the pool of immature bases. When a base was lost due to dilution, we selected a base at random with equal probability and removed it from the pool of mature or immature bases depending on whether the chosen base was mature or immature. When a base is removed, all the needle proteins associated with it were also lost from the system. Only immature bases are capable of exporting ruler and needle proteins. Once the needle length achieves a length is greater than or equal to the maturation length, $l_R$, the next ruler protein checking event makes the base mature and it is removed from the pool of immature bases and added to the pool of immature bases.

In our model needle protein export always increases the number of needle proteins associated with a given base. In doing so, we ignored a number of possible scenarios. For example, a needle protein could be exported and simply not bind to the base and could be lost in the extracellular space. Alternately, a needle protein that is present near the tip of the the needle could also unbind. Some of these free proteins available in the extracellular space could re-bind to the needle, although the extracellular volume would be so large compared to volume of the cell that the chances of a re-binding event happening would be virtually zero. Nevertheless, the export events mentioned above would only change the binding rate of the needle proteins in our ODE model, $\beta_O$, to some new effective binding rate $\beta'_O$. This would only change the numerical relationship between results of our ODE model and inputs to our statistical model and would not have any impact on the general behaviour of the system.

We implemented the stochastic simulation using the standard Gillespie-Doob approach [2]. The rates of the reactions were calculated using the parameters defined in the deterministic model: $Q$'s, $\beta$'s, and $\lambda$'s. The rates of the individual reactions are specified on the arrows in figure 1. All simulations were executed till the system achieved a steady state.

### Error-prone ruler

To incorporate the uncertainty in measurement due to conformational changes in the ruler protein, we represent the maturation probability using a logistic function;

$$F(L; l_R, s) = \frac{1}{1 + e^{-\left(\frac{L - l_R}{s}\right)}}. \tag{16}$$

where $L$ : the needle length, $l_R$ : the length of the exact ruler, and $s$ : the uncertainty in measurement. Figure 2 shows the difference in the maturation probabilities between the exact and logistic rulers.

We implemented the error-prone ruler in our stochastic simulations as follows. Instead of having a ruler of a fixed length $l_R$, we randomly chose a ruler from a population of lengths that obeys the logistic

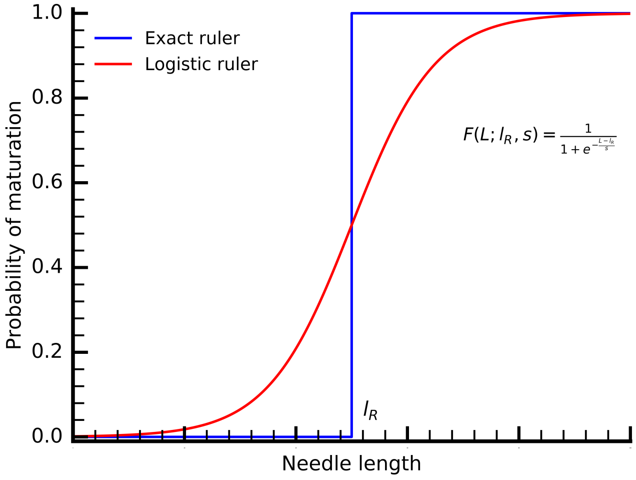


Figure 2: Maturation probabilities using exact and logistic rulers.

distribution parameterized with $l_R$ as the mean and $s$ as the uncertainty. Thus maturation now occurs for this new ruler length which is different for each independent ruler protein exported. Note that this only affects maturation, and all other reactions of the system occur as before. It is unclear whether the measurement uncertainty that arises due to conformational differences in ruler protein would remain fixed for rulers of different lengths or vary with the ruler length itself. We explored both of these possibilities in our simulations by keeping $s$ fixed and increasing its value proportionally with $l_R$ ($s =$ $0.1\ l_R$, $0.2\ l_R$). Our analysis suggests that an error-prone ruler offers a robust control over the length regulation for both the cases, when $s$ is fixed and when $s$ is increased relative to $l_R$.
